# Effects of Spatial Constraints of Inhibitory Connectivity on the Dynamical Development of Criticality in Spiking Networks

**DOI:** 10.1101/2024.12.04.626902

**Authors:** Felix Benjamin Kern, Takahisa Date, Zenas C. Chao

## Abstract

Neural systems are hypothesized to operate near criticality, enhancing their capacity for optimal information processing, transmission and storage capabilities. Criticality has typically been studied in spiking neural networks and related systems organized in random or full connectivity, with the balance of excitation and inhibition being a key determinant of the critical point of the system. However, given that neurons in the brain are spatially distributed, with their distances significantly influencing connectivity and signal timing, it is unclear how the spatial organization of excitatory and inhibitory connectivity influences the network’s self-organization towards criticality. Here, we systematically constrain the distance and density of inhibitory connectivity in two-dimensional spiking networks and allow synaptic weights to self-organize with activity-dependent excitatory and inhibitory plasticity in the presence of a low level of stochastic intrinsic activity. We then investigate the relationship between inhibitory connectivity, synaptic weights, and the resulting network activity during and after development. We find that networks with longer-range inhibitory synapses tend towards more supercritical behavior compared to networks with a similar number of shorter-range inhibitory synapses. We show that this distance dependence is a consequence of weaker long-range synapses after development due to the presence of synaptic delays, which shift most spike pairs outside of the potentiation window of the inhibitory learning rule.

## Introduction

The brain criticality hypothesis states that neurons in the brain are organized in such a way that their collective activity lies at or near the phase transition between ordered and random activity (Beggs 2007; Beggs and Timme 2012). At this critical point, neural activity is characterized by self-similarity and scale-free activity patterns. Criticality in neural networks has been shown to be beneficial in many ways. Criticality leads to optimal computational capacities (Legenstein and Maass 2007; Bertschinger and Natschläger 2004) and optimal information transmission (Beggs and Plenz 2003; Shew et al. 2011). Networks operating at or near criticality exhibit high information storage capacity (Beggs and Plenz 2004; Haldeman and Beggs 2005; Shew et al. 2011) and achieve their maximum dynamic range, making them highly sensitive to changes in external stimulation (Kinouchi and Copelli 2006; Shew et al. 2009).

Self-organized criticality, referring to systems that intrinsically evolve towards a critical point, was first discussed in the field of physics with the sandpile model (Bak, Tang, and Wiesenfeld 1987). There, avalanches are triggered in a scale-free manner, meaning that there is a power-law relationship between the size of avalanches and their frequency or probability of occurrence. Scale-free neural activity, so-called neuronal avalanches, were subsequently discovered in cortical neurons as a signature of self-organized criticality (Beggs and Plenz 2003). Neuronal avalanches have been observed in various in-vitro and in-vivo experimental models (Beggs and Plenz 2003; Tetzlaff et al. 2010; Ponce-Alvarez et al. 2018).

How the brain develops into the critical state is an active research question (Tetzlaff et al. 2010; Hesse and Gross 2014; Plenz et al. 2021). Previous research has revealed that network structure, long-term synaptic plasticity rules, and homeostatic mechanisms can drive networks to develop into a critical state (Zeraati, Priesemann, and Levina 2021; Stepp, Plenz, and Srinivasa 2015; Del Papa, Priesemann, and Triesch 2017; Rubinov et al. 2011). Meanwhile, short-term plasticity is known to expand the critical regime from a finely balanced point to a broad region in parameter space (Zeraati, Priesemann, and Levina 2021). Finally, regardless of how criticality is established, the single most important parameter for its maintenance is the balance of excitation and inhibition (E/I balance) within a network (Poil et al. 2012; Mazzoni et al. 2007; Benayoun et al. 2010).

Most research into criticality has been conducted in randomly or fully connected networks, ignoring the spatial dimensionality of the brain and the resulting structural anisotropies. The effect of spatial constraints has been investigated in the context of excitatory connectivity (Wilkerson and Moschoyiannis 2021; Poil et al. 2012), while synaptic delays have been shown to affect how quickly networks settle into a steady state (Larremore et al. 2011). However, to our knowledge, there has been no work investigating the impact of inhibitory connectivity or of the influence of conduction delays on the steady-state activity itself.

Here, we investigate the effect of spatial constraints in inhibitory connectivity on the long-term development of criticality, marking a first foray into a more comprehensive treatment of criticality in networks with spatial dimensions. In addition, unlike related studies using spatial constraints for connectivity only (Wilkerson and Moschoyiannis 2021; Poil et al. 2012), we explicitly model the delays imposed by conduction along axons and find that such delays have a profound effect on the development of synaptic weights.

## Methods

### Neuron model

We chose a basic leaky integrate-and-fire neuron model with a membrane potential *u* evolving according to

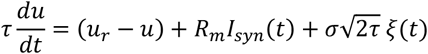

with membrane time constant τ = 30 ms, membrane resistance *R*_*m*_ = 100 MΩ, and resting membrane potential *u*_*r*_ = − 69 mV. When a neuron’s membrane potential reached the firing threshold *θ* = − 54 mV, a spike was emitted, and the membrane potential was clamped to *u* = *u*_0_ = −74 mV for 3 ms (excitatory neurons) or 2 ms (inhibitory neurons). Noise was injected into the membrane potential in the form of an Ornstein-Uhlenbeck process with a Gaussian random variable ξ(*t*) and a standard deviation *σ*=5 mV, yielding an intrinsic firing rate of approximately 0.28 s^−1^. A sample membrane potential trace of an isolated excitatory neuron is shown in Figure 1A.

**Figure 1.**
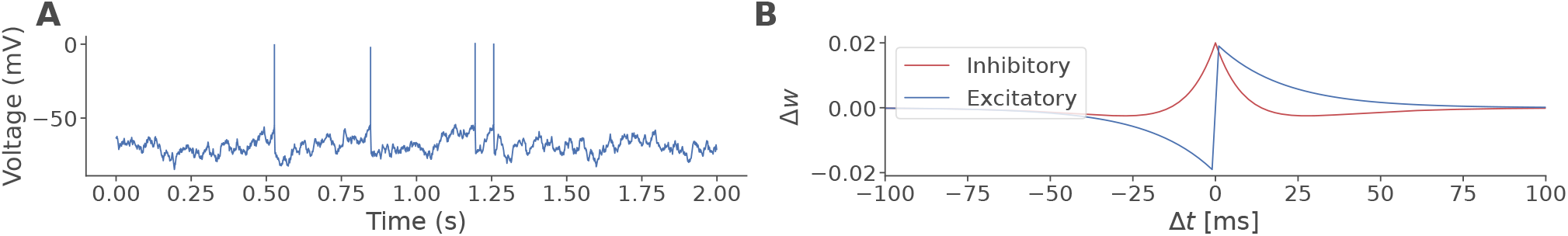
Model dynamics. **(A)** Sample neuronal voltage trace of a disconnected excitatory neuron. **(B)** Excitatory and inhibitory STDP rules, with Δ*t* = *t*_*post*_ − *t*_*pre*_.

### Synapse model

Synaptic currents were modeled in a conductance-based manner, following

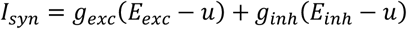

with excitatory reversal potential *E*_*exc*_ = 0 mV and inhibitory reversal potential *E*_*inh*_ = −100 mV. Synaptic conductances evolved according to

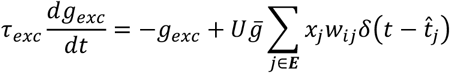

and

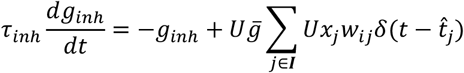

for excitatory and inhibitory synapses, respectively, with excitatory time constant τ_*exc*_ = 2 ms, inhibitory time constant τ_*inh*_ = 4 ms, scale factor 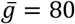 nS, the sets of presynaptic excitatory and inhibitory neurons ***E*** and ***I***, respectively, synaptic weight *w*_*ij*_, the Dirac delta function *δ*(·) and presynaptic spike times 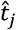.

Synapses were subject to short-term depression, using a deterministic simplification of the model in (Tsodyks and Markram 1997). We modeled the depression variable *x*_*j*_ as a property of the presynaptic neuron *j*,

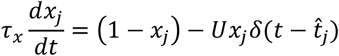

with recovery time constant τ_*x*_ = 150 ms and release fraction *U* = 0.4.

Finally, all synapses were also governed by long-term spike-timing dependent plasticity (STDP). In excitatory synapses, we used standard STDP (Song, Miller, and Abbott 2000), with updates to the dimensionless weight value *w*_*ij*_ ∈ [0,1] of the excitatory synapse from neuron *j* to neuron *i* following

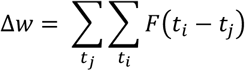

where

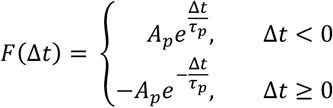

with τ_*p*_ = 20 ms and *A*_*p*_ = 0.01.

For inhibitory synapses, we employed a symmetrical Hebbian STDP rule (Ikeda, Akita, and Takahashi 2023; Stepp, Plenz, and Srinivasa 2015) whose weight update function follows

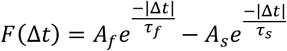

where *A*_*f*_ = 0.02, *A*_*s*_ = 0.01, and the time constants τ_*f*_ = 10 ms and τ_*s*_ = 20 ms. This rule gives rise to a “Mexican hat” shaped weight update function resembling the homeostatic rule in Vogels et al. (2011). Both excitatory and inhibitory rules are illustrated in Figure 1B.

Spikes were delivered to synapses with a distance-dependent axonal delay corresponding to a conduction velocity of 0.15 m/s along a direct linear path from presynaptic to postsynaptic neuron location.

### Network structure

We investigated networks of *n*_*exc*_ = 80 excitatory and *n*_*inh*_ = 20 inhibitory neurons scattered randomly across a two-dimensional disk with a radius of 2 mm. Network structure was established based on the spatial proximity of neurons, as illustrated in Figure 2A. Specifically, for a given presynaptic neuron of type *k* ∈ {*exc, inh*}, all neurons located within a radius of *r*_*k*_ were considered potential postsynaptic targets. From among this subset, synaptic connections were established at random with a uniform probability *p*_*k*_. While excitatory connectivity was fixed at *r*_*exc*_ = 1.3 mm and *p*_*exc*_ = 0.6, we varied inhibitory connectivity systematically, with *r*_*inh*_ ∈ {0.5, 1,2,3,4} mm and *p*_*inh*_ ∈ {0.1, 0.3, 0.5, 0.7, 1}, yielding a 5-by-5 grid of parameter settings to explore.

**Figure 2.**
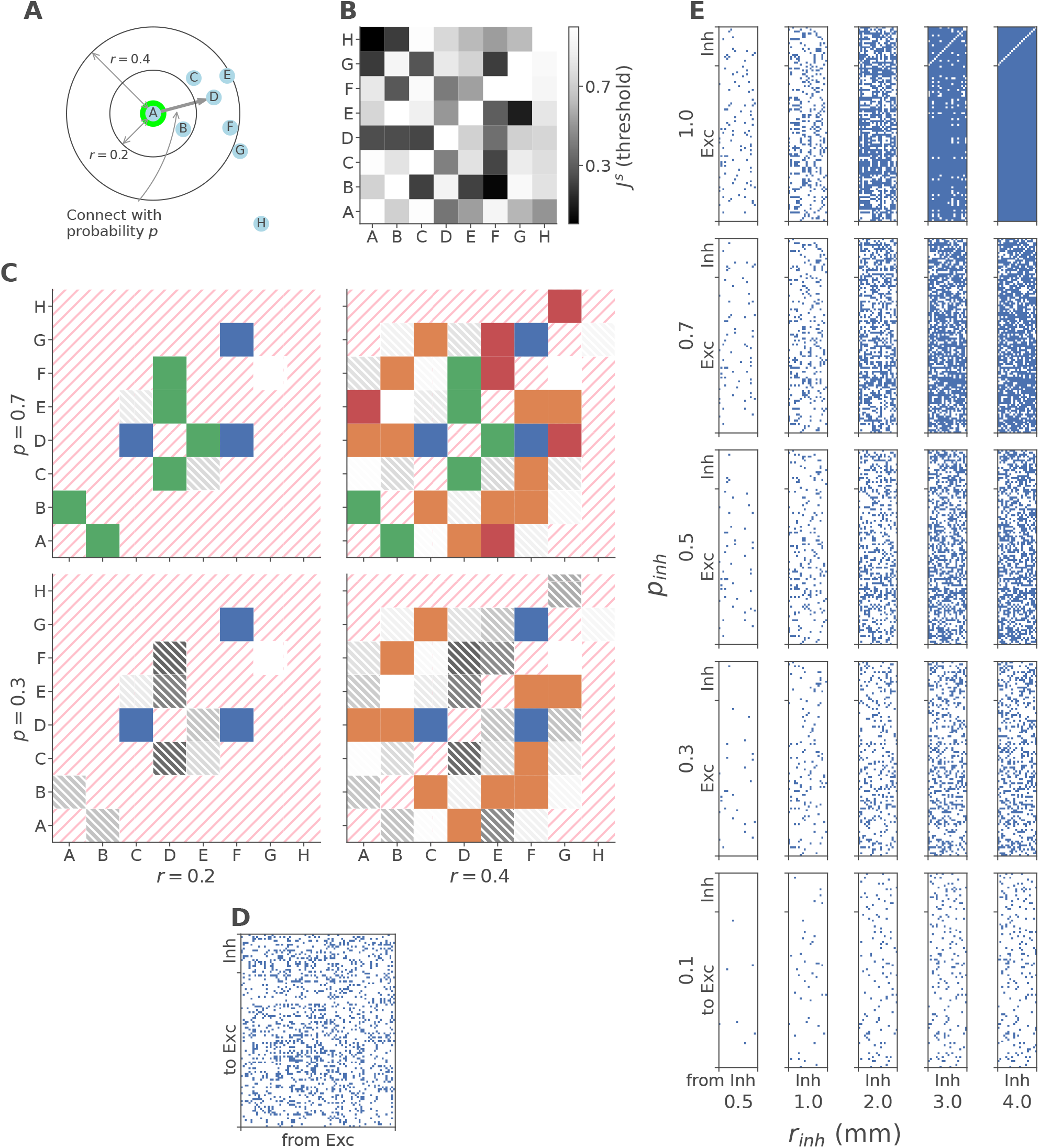
Illustration of the additive process used to generate connectivity. **(A)** Locations of the eight neurons (A-H) used in this illustration. Black circles are drawn at radius *r* = 0. 2 and *r* = 0.4 around the highlighted neuron A for reference. **(B)** The adjacency matrix scaffold ***J***^*s*^ for this illustration, drawn pseudorandomly from a uniform distribution. Darker squares will be connected at lower probability thresholds *p*. **(C)** The resulting adjacency matrices for two values each of *r* and *p*. Neuron pairs that are too distant (i.e., their distance is greater than *r*) are disconnected, along with the diagonal (i.e., no autapses), as illustrated by the red hatching. Neuron pairs that are within the spatial window for *r*, but whose value of *J*^*s*^ exceeds *p*, are disconnected and shown hatched in grey (same color scale as in panel B). Neuron pairs that meet the criteria for both distance (*< r*) and probability (*J*^*s*^ < *p*) are connected and shown in solid color, with different hues indicating connections that are present throughout (blue), added by increasing *r* (green), increasing *p* (orange), or increasing both *r* and *p* together (red). **(D-E)** Adjacency matrices of one network “prototype” (see text). Blue pixels indicate existing connections. **(D)** Excitatory connectivity, drawn from *r*_*exc*_ = 1.3 mm and *p*_*exc*_ = 0.6. **(E)** Inhibitory connectivity across the 5×5 grid of values for *r*_*inh*_ and *p*_*inh*_.

To minimize the impact of structural stochasticity, we opted to maintain as much of the structure as possible across parameter settings. Specifically, spatial positions and excitatory connections were maintained across parameter settings, while the inhibitory connections were constructed in an additive fashion with increasing *r*_*inh*_ and *p*_*inh*_. To achieve this, we initialized a scaffold 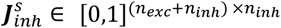 of the adjacency matrix ***J***_*inh*_ with a single set of pseudo-random numbers for all 25 parameter settings. Then, we selected connections using the specific values of *r*_*inh*_ and *p*_*inh*_, defining the elements of ***J***_*inh*_ as

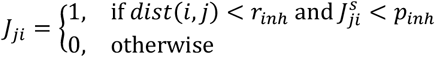

In this way, the set of inhibitory connections in each network is a subset of the connections in all networks with greater or equal *r*_*inh*_ and *p*_*inh*_. The principle of this approach is illustrated with a toy example with eight neurons in Figure 2. Figure 2A shows the spatial locations of the neurons, Figure 2B a randomly drawn adjacency matrix scaffold ***J***^*s*^, and Figure 2C shows the resulting adjacency matrices for two values of *r* and *p* each.

We use the term “network prototype” to refer to a single set of 25 networks created in this way, each sharing in common the neuron locations, excitatory connectivity, and parts of inhibitory connectivity. We independently generated and simulated a total of 10 network prototypes for analysis. The adjacency matrices of one of these are presented in Figure 2D, showing the excitatory portion, which is identical across parameter settings, and in Figure 2E, showing the inhibitory portions across the 5×5 grid of *r*_*inh*_ and *p*_*inh*_.

### Development

While the structure of the networks, i.e., the presence or absence of synapses, was fixed, we initialized all synaptic weights to zero to imitate an immature neuronal network. We then simulated each network for 15 hours, allowing weights to develop according to the plasticity rules outlined above. To account for the influence of stochasticity in intrinsic neuronal activity, each simulation was repeated five times with different random seeds.

### Criticality

To investigate criticality properties of the developing networks, we analyzed the neural avalanches and calculated the criticality variable Δ*p* as follows. First, we counted spikes across the network in bins of 12 ms duration. Contiguous sections of non-zero spike counts were considered “avalanches” with a size *s* corresponding to the total number of spikes fired. In temporal windows of 5 minutes, we then calculated the probability *p*(*s*) of seeing an avalanche of a particular size and approximated the resulting distribution with a power law function, *p*(*s*) = *a s*^*b*^, using linear least-squares fitting. Finally, the desired measure Δ*p* was defined as the average deviation of the data from this fit across all avalanche sizes, i.e.,

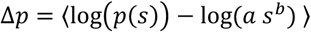

## Results

Our results are presented as follows. First, we will briefly cover the development of the networks over 15 hours of simulation time, giving an intuition for the steady-state activity patterns and their emergence. Then, we will focus on the steady state, comparing activity patterns across different parameter settings and confirming that our outcomes broadly agree with previous findings on the boundary conditions for criticality. Finally, we will investigate how the spatial constraint on inhibitory connections affects criticality, showing that networks with more long-range inhibitory connections tend to be more supercritical than networks with more short-range connections, since long-range inhibitory synapses are weaker than short-range synapses and therefore less able to counter excitatory influence.

### Development

Starting from an initial state with connections established but ineffective (w=0), networks followed a largely conserved trajectory of changes in weights and activity irrespective of the constraints on inhibitory connectivity. Figure 3 shows an example, elaborated below, of this trajectory in one network. Other runs (i.e., simulations of the same network with different random seeds for intrinsic activity) and other networks and network prototypes followed a qualitatively very similar profile.

**Figure 3.**
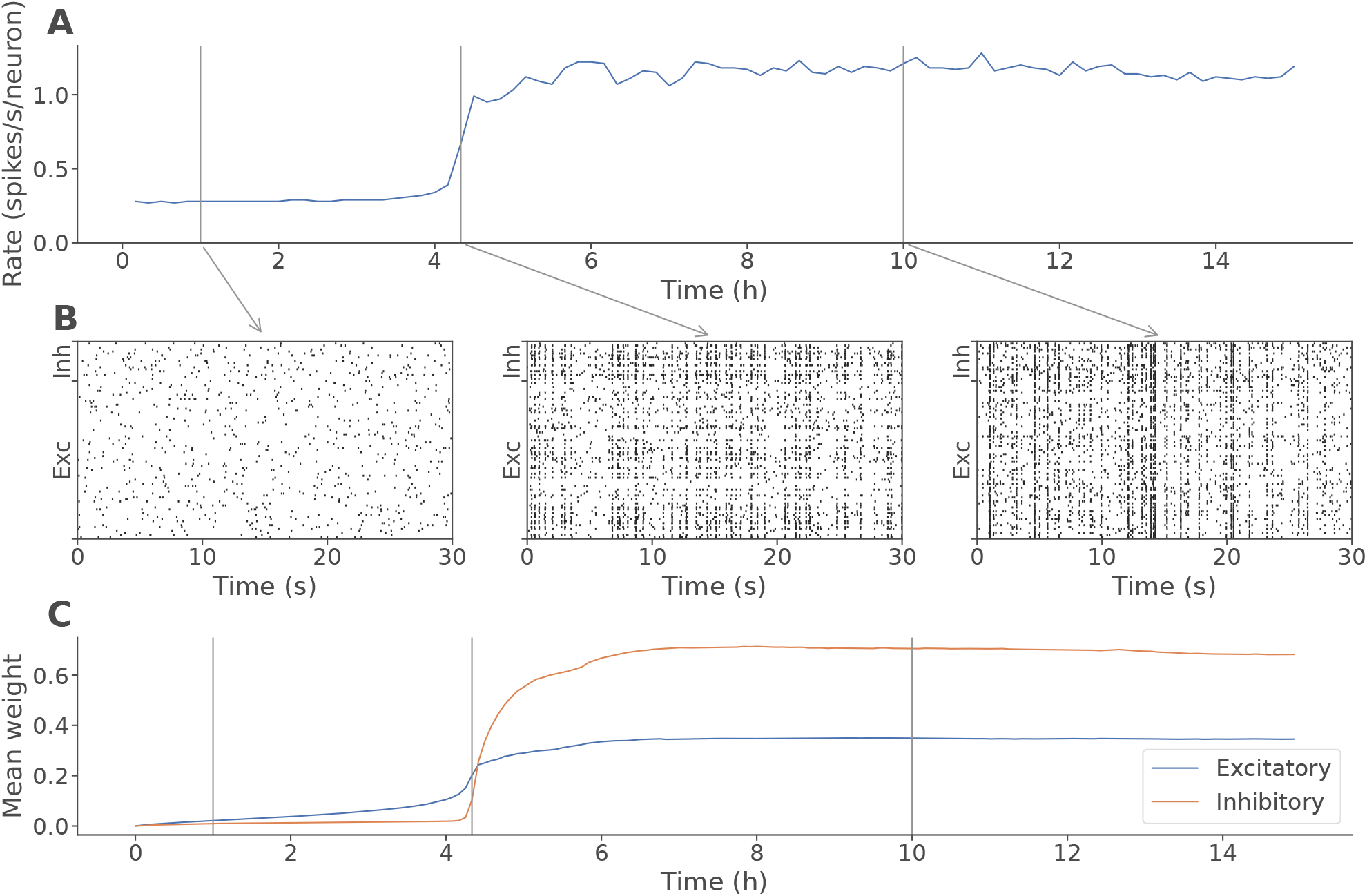
Development of a sample network (*r*_*inh*_ = 4 mm, *p*_*inh*_ = 0. 7). **(A)** Mean firing rate in spikes per second per neuron. **(B)** Raster plots of network activity during 30 second samples taken from the time points indicated in panel A (1h, 4h20min, 10h). **(C)** Mean synaptic weight across existing excitatory connections (blue) and inhibitory connections (orange).

Initial activity was sparse, uncoordinated, and driven exclusively by intrinsic voltage fluctuations (Figures 3A and 3B). After four to five hours, excitatory weights (Figure 3C) had grown sufficiently strong to cause coordinated activity, which quickly escalated to highly synchronized network-wide bursting (Figure 3B, middle). This, in turn, drove inhibitory weights to catch up, providing a degree of counterbalance and allowing networks to control activity into an approximately steady state within one to two hours (Figure 3B, right). Both firing rate and weights then remained approximately constant until the end of the 15-hour simulation period.

### Criticality after development

To assess the activity patterns at steady state, we focused on the final two hours of simulation (13-15 h) in each network and run. Figure 4A shows raster plots of one network prototype, taken from the final 20 seconds in each of the 25 settings of *r*_*inh*_ and *p*_*inh*_. In Figure 4B, we illustrate the derivation of the criticality measure Δ*p* on data from three of these networks, exhibiting supercritical behavior (left, *r*_*inh*_ = 0.5 mm and *p*_*inh*_ = 0.1, notice the large number of large avalanches), approximately critical behavior (center, *r*_*inh*_ = 2 mm and *p*_*inh*_ = 0.5), and subcritical behavior (right, *r*_*inh*_ = 4 mm and *p*_*inh*_ = 1, with few large avalanches).

**Figure 4.**
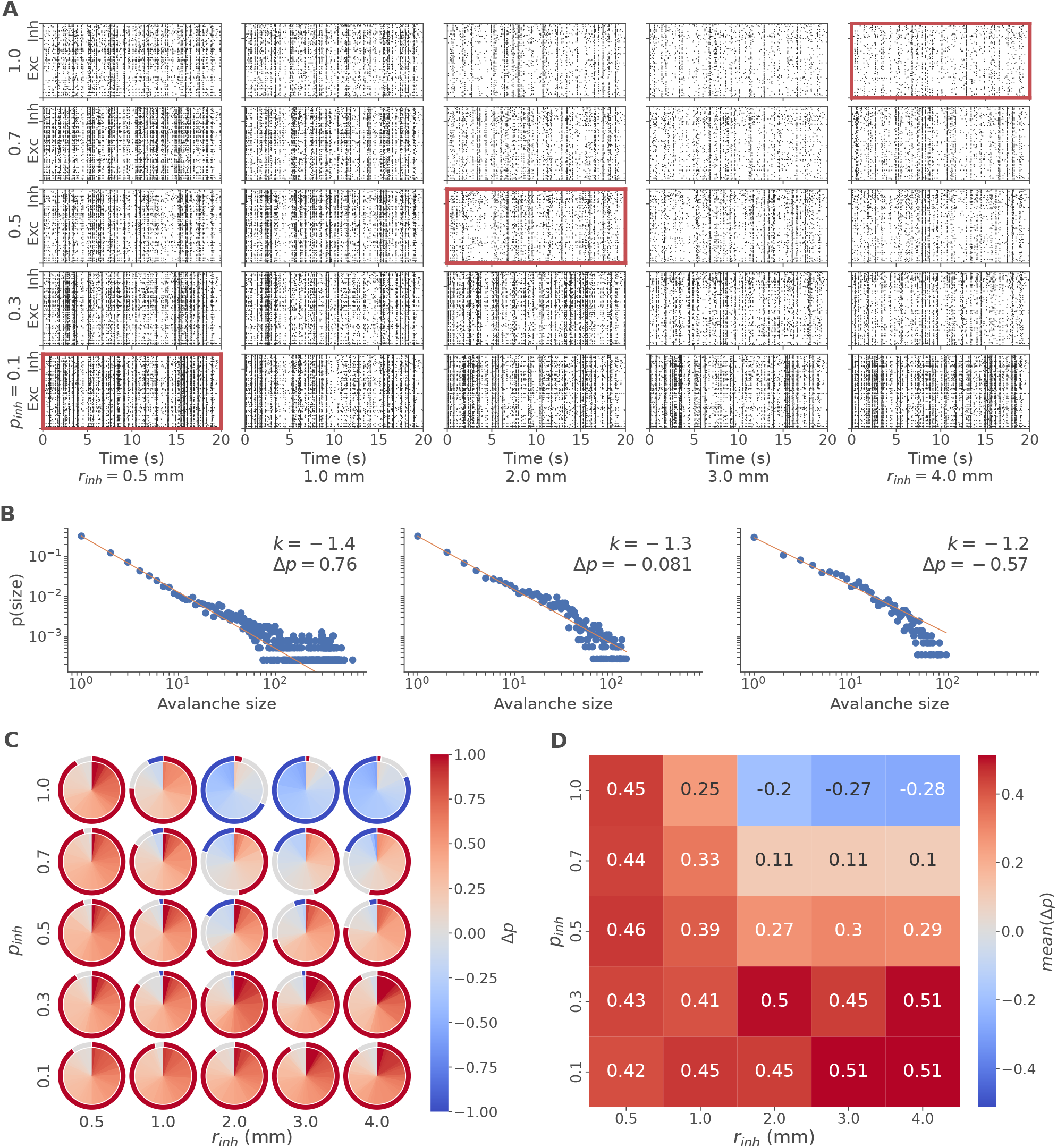
Criticality after development. **(A)** Spike raster plots showing the firing patterns of a sample network prototype across the grid of *r*_*inh*_ and *p*_*inh*_ after 15 hours of development. **(B)** Frequency of avalanches plotted against their respective size (points) along with the corresponding power-law fit (solid line), illustrating the calculation of *Δp*, which is the mean error between the observed frequencies and the power-law fit. The three plots correspond to the highlighted raster plots in panel A and summarize the final 5 minutes of activity in each of these networks. Inset: Slope *k* and criticality outcome *Δp*. **(C)** Distribution of *Δp* across the grid of *r*_*inh*_ and *p*_*inh*_. *Δp* was calculated in each run of each network from activity between 13 and 15 hours. The inner circle represents all 5 runs x 10 networks in each grid cell as 50 equally sized wedges, color-coded and sorted by *Δp*. A small number of outliers with *Δp* > 1 are color-coded as dark red. The outer ring represents the fraction of supercritical (red), critical (|*Δp*| < 0.1, grey), and subcritical (blue) values. **(D)** The mean of *Δp* across all runs and networks in each grid cell.

The average values of Δ*p* in the final two hours for each of the 25 parameter configurations are shown individually in Figure 4C. The mean of Δ*p* across networks and runs, indicative of the expected outcome for each setting of *r*_*inh*_ and *p*_*inh*_, is shown in Figure 4D, and the associated standard deviation in Supplementary Figure 1. We observed supercritical behavior throughout most of the parameter space, being the majority outcome in nearly all conditions, except where inhibitory connectivity was very dense with *r*_*inh*_ ≥ 2 mm and *p*_*inh*_ = 1. Criticality, which we define as |Δ*p*| < 0.1, was seen to emerge infrequently in all parameter settings, but was most common at intermediate inhibitory connectivity densities (*r*_*inh*_ ≥ 2 mm, *p*_*inh*_ ∈ {0.5, 0.7}), while at the highest densities (*r*_*inh*_ ≥ 2 mm, *p*_*inh*_ = 1), subcritical behavior predominated. In sum, the higher the density of inhibitory connections at the outset, the lower the value of Δ*p* became after development.

### Excitation-inhibition balance in the developed state

This is not prima facie surprising: Previous work (Poil et al. 2012) has shown that criticality typically emerges when excitation and inhibition hold each other in balance. This is hinted at in the results presented above, where we have used the initial constraint. To verify the role of E/I balance in maintaining criticality, we analyzed the steady state of the networks following development. After development, most synapses of all types had weight values approaching either 0 or 1 (Supplementary Figure 2). Therefore, we set a weight threshold of 0.5, considering any synapses at or above the threshold as active, and any below as inactive. Then, we counted the number of active excitatory and inhibitory synapses onto each neuron (the effective in-degree *k*^*eff*^) and averaged across neurons. Plotting the inhibitory against the excitatory effective degree of all 1250 networks (25 parameter settings, 10 networks, 5 runs), we see a clear linear separation between supercritical networks, whose inhibitory effective degree is low, and subcritical networks, whose inhibitory effective degree is high relative to their excitatory effective degree (Figure 5A). Extracting just the critical networks (Figure 5B), we find that a linear relationship fits the data well, with 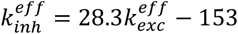 and a Pearson correlation coefficient of 0.69 (p = 8.6e-27, n = 179 critical networks).

**Figure 5.**
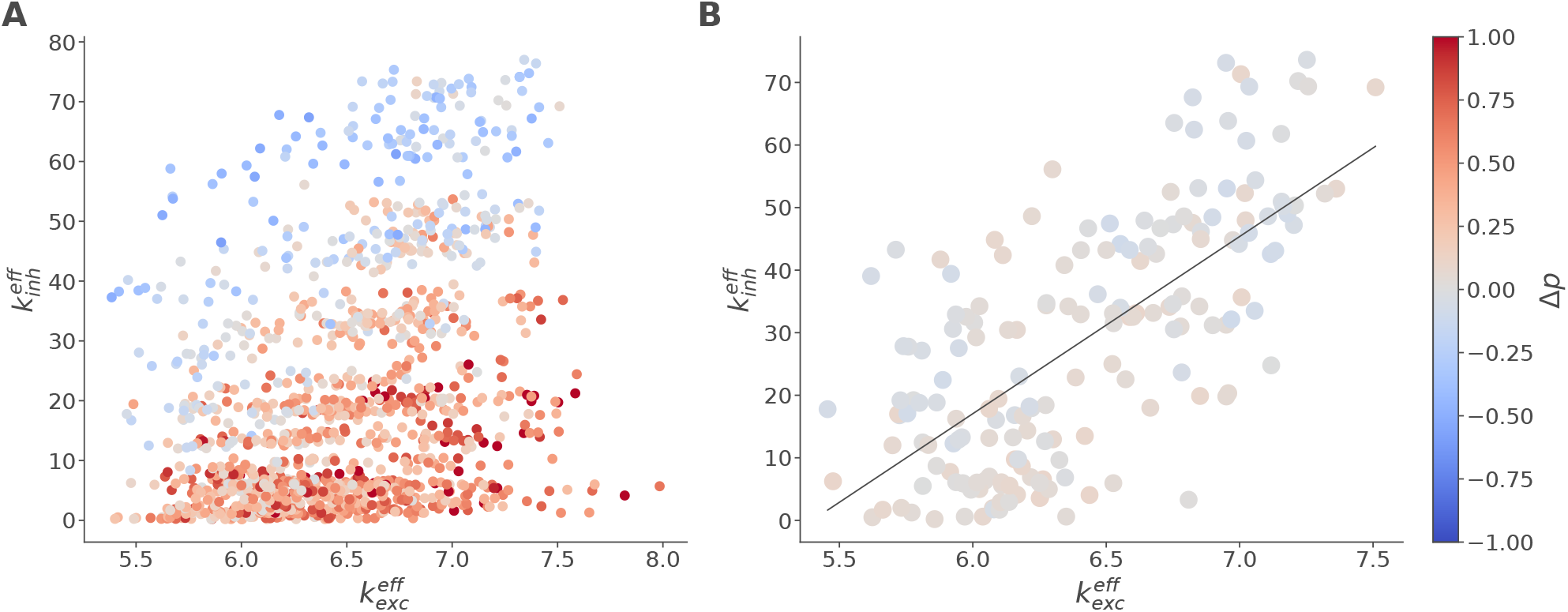
Balance of excitation and inhibition is tightly linked to criticality. **(A)** Network degree, i.e., the mean number of incoming connections with *w* ≥ 0.5 per neuron, separated by the type of incoming connection and plotted against each other. Each dot represents one network run in its weight configuration after 15 hours of development. Color represents *Δp*. **(B)** The same data in the subset of networks with |*Δp*| < 0.1. The black line is the linear least-squares fit to these data points with slope 28 and intercept -153.

### Effect of spatial constraint

Unlike most previous research, our approach to manipulating networks not only affects connection density or weight, but also network topology based on the spatial relationships between neurons. That is, we wanted to know how the length of inhibitory connections affects criticality independently of connection density. However, the parameters we chose to manipulate the networks, *r*_*inh*_ and *p*_*inh*_, both affect connection density. Therefore, to cleanly separate the effects of connection length from those of connection density, we calculated the average inhibitory in-degree and the average inhibitory connection length for each network, sliced the resulting space into bins of roughly equal inhibitory degree, and analyzed the effect of connection length separately in each slice as detailed below. Note that, unlike in the above analysis on E/I balance, here we used the connection blueprint, i.e., all existing connections regardless of their weight after development, to quantify the structurally defined connectivity and thereby capture the effect of the initial constraint.

### Long-range inhibitory networks yield larger avalanches

First, we mapped all networks into the space of mean inhibitory degree *k*_*inh*_ (Figure 6A) and mean inhibitory connection length *l*_*inh*_ (Figure 6B). There is clear clustering of networks, an artifact of the discrete set of parameter values. Plotting *l*_*inh*_ against *k*_*inh*_ (Figure 6C), we also see an obvious absence of any networks in the lower right half of the plot (high degree with short average length) due to the spatial nature of our model. Considering the dynamics, we see that networks with greater *k*_*inh*_ clearly show lower *Δp* (Pearson’s r = -0.59, p = 4e-120, n = 1250; Figure 6A), corroborating the above findings on E/I balance even when ineffective connections are included.

**Figure 6.**
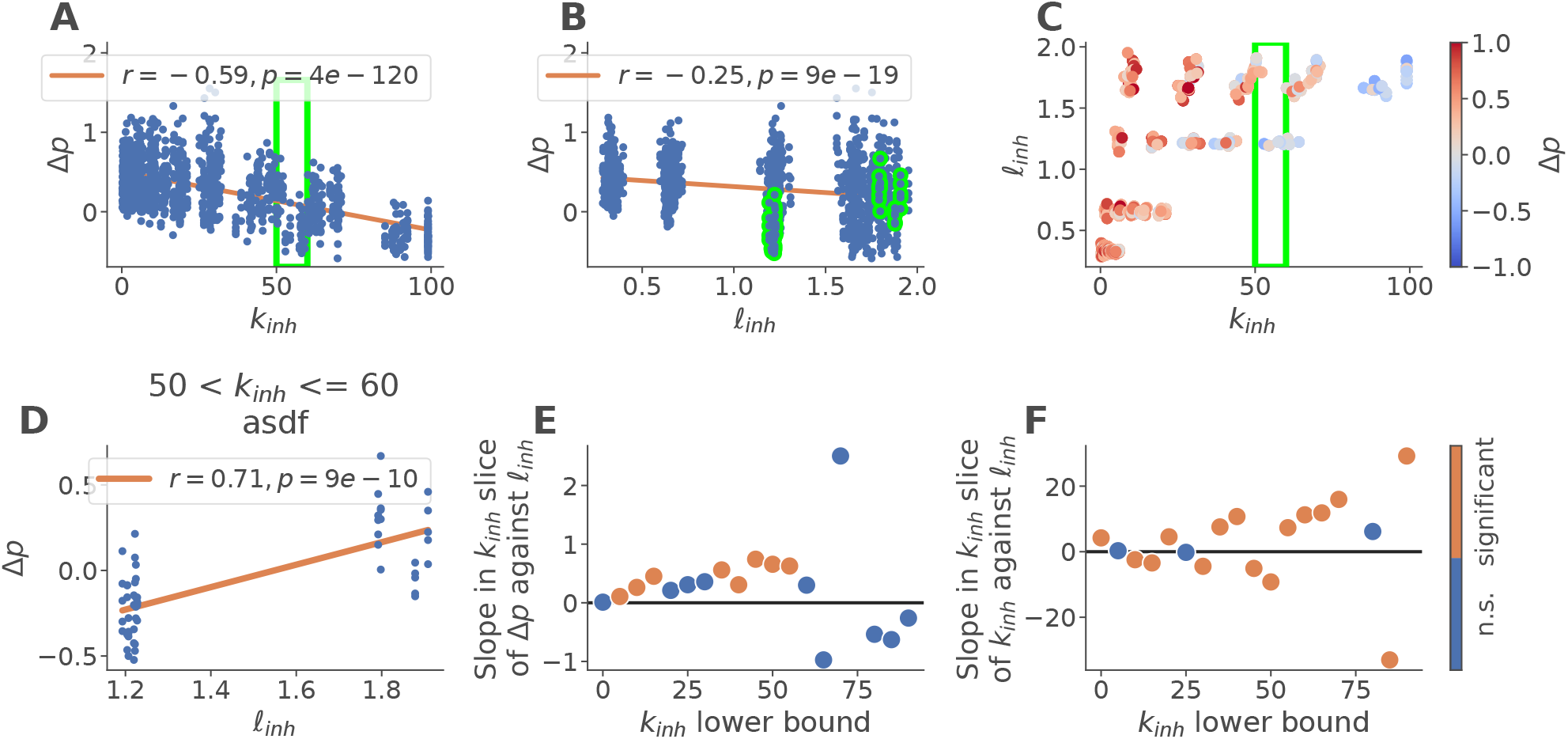
Long-range inhibitory networks yield larger avalanches. **(A)** Criticality outcome (*Δp*) plotted against the mean inhibitory degree (structurally defined *k*_*inh*_, i.e., including all existing connections regardless of weight) along with a linear fit (orange). The green box highlights one slice of *k*_*inh*_ containing networks of approximately equal degree referred to in other panels. **(B)** Criticality outcome plotted against mean length of inhibitory connections (structurally defined *l*_*inh*_) along with a linear fit. Points highlighted in green correspond to the slice in panel A. **(C)** Criticality outcome (*Δp*, point color) shown as a function of *k*_*inh*_ and *l*_*inh*_. **(D)** Sample correlation between *l*_*inh*_ and *Δp* in the slice indicated in panels A-C. The orange line indicates the best-fit linear model. **(E)** Slopes of the linear fit relating *l*_*inh*_ to *Δp* in each slice of *k*_*inh*_. Orange color indicates significant linear correlations (Bonferroni-corrected *p* < 0.05). **(F)** Slopes of the linear fit relating *l*_*inh*_ to *k*_*inh*_ in each slice of *k*_*inh*_. Orange color indicates significant linear correlations (Bonferroni-corrected *p* < 0.05).

The relationship between Δ*p* and *l*_*inh*_ is similar, but weaker (r = -0.19, p = 9e-19, n = 1250). Noting the dependence between degree and connection length (Figure 6C), we sought to isolate the influence of connection length *l*_*inh*_. To do so, we sliced the data into overlapping bins of width 10 along the *k*_*inh*_ axis, such that each slice contained networks of approximately equal degree but potentially very different connection lengths. We then correlated *l*_*inh*_ in these slices with Δ*p* (see an example in Figure 6D). We see that greater *l*_*inh*_ predicts greater Δ*p* across a substantial portion of the slices (Figure 6E and Supplementary Figure 3). In other words, networks with longer inhibitory connections are more supercritical than networks with a similar number of shorter inhibitory connections. Indeed, greater *l*_*inh*_ could even tip networks from subcritical to supercritical states, as in the example in Figure 6D.

We note that the slice-wise positive correlations do not conflict with the overall negative correlation (Figure 6B), but rather show that the overall correlation result is largely due to the confound between *l*_*inh*_ and *k*_*inh*_. In addition, as we show in Figure 6F and Supplementary Figure 4, the correlation between *l*_*inh*_ and Δ*p* in slices is not mediated by variations in *k*_*inh*_ within the slices, which are not related to *l*_*inh*_ in a systematic manner.

### Long-range inhibitory connections are less functional

Next, we focused on possible reasons for this relationship. The mediating factor between initial connectivity and critical dynamics after development is, of course, synaptic weight. All else being equal, we should expect the E/I balance of weights to constitute a key driver of criticality. Indeed, as shown in Figure 7A, the E/I weight ratio, i.e., the ratio of mean excitatory to mean inhibitory synaptic weight, was positively correlated with Δ*p* in most slices of similar *k*_*inh*_, corroborating the related result for the E/I balance of effective degree in Figure 5. In turn, we find that E/I weight ratio is largely determined by *l*_*inh*_ (Figure 7B), with networks with longer average connection length showing a larger E/I weight ratio. This effect is driven almost entirely by smaller inhibitory weights (Figure 7C) rather than by larger excitatory weights (Figure 7D).

**Figure 7.**
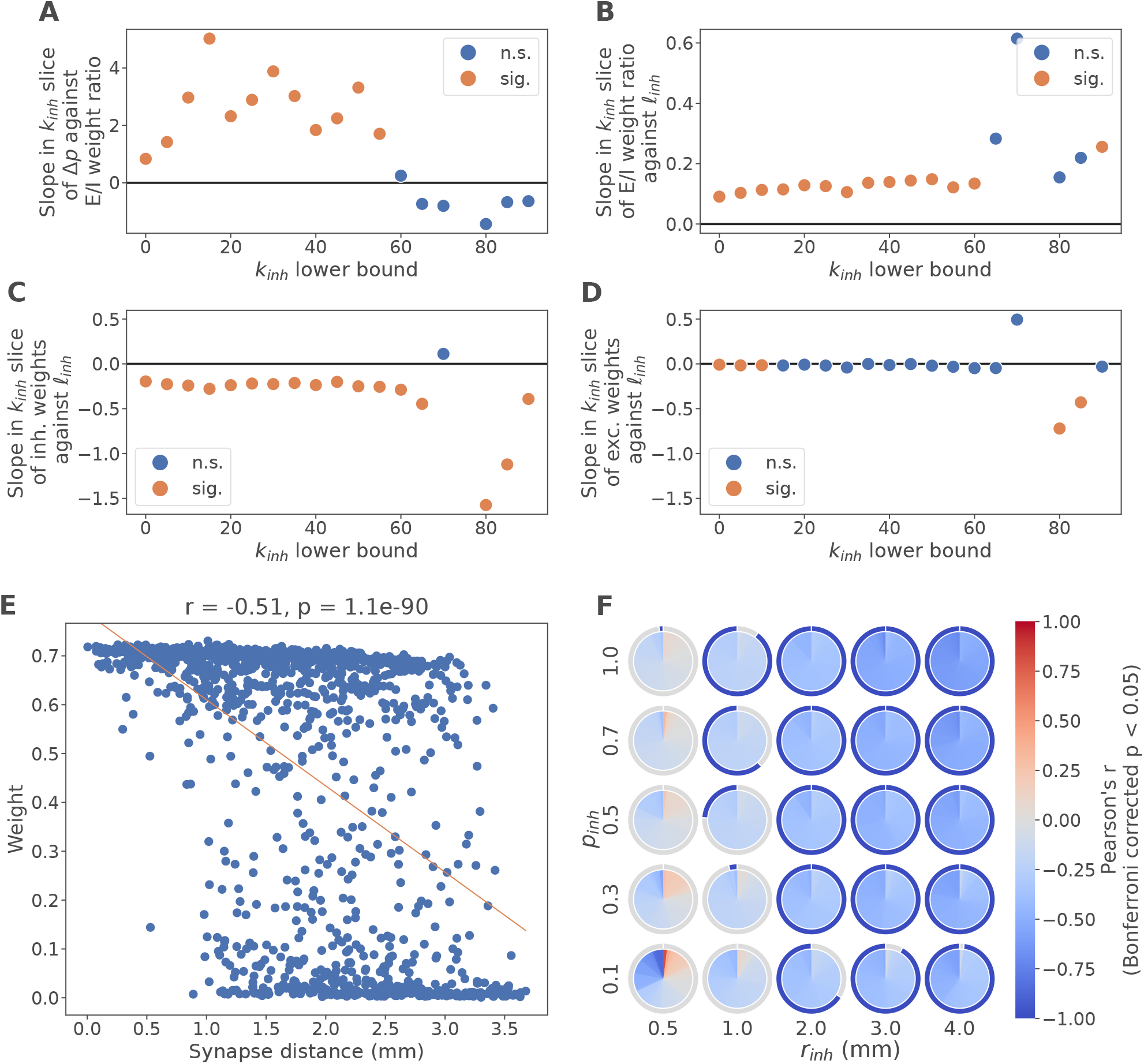
Long-range inhibitory connections are less functional. **(A-D)** Slopes of linear fits across networks in slices of *k*_*inh*_ (see Figure 6 for an explanation of the slices, and Supplementary Figures 5-8 for scatter plots of each slice). Orange color indicates significant linear correlations (Bonferroni-corrected *p* < 0.05). The linear fits relate the following pairs of values: **(A)** The criticality outcome, *Δp*, to the ratio of mean excitatory and mean inhibitory weights (E/I weight ratio). **(B)** The E/I weight ratio to the mean inhibitory synaptic distance across neurons, *l*_*inh*_. **(C)** The mean weight of inhibitory connections to *l*_*inh*_. **(D)** The mean weight of excitatory connections to *l*_*inh*_. **(E-F)** Inhibitory weight and synaptic distance correlated within networks. **(E)** Weight of inhibitory synapses (mean across development) plotted against the distance between the corresponding neurons in a representative sample network (same as in Figure 3; *k*_*inh*_ = 68). The orange line represents the linear fit. **(F)** Distributions of Pearson’s *r* of the above correlation within networks (i.e., between weight and distance of inhibitory synapses). The inner circle represents all 50 networks and evaluations in each grid cell as equally sized wedges, color-coded and sorted first by significance, then by *r*. The outer ring indicates significant negative correlations (blue) and insignificant correlations (grey); no significant positive correlations were found. Significance was assessed with a two-tailed t-test and Bonferroni correction at a level of *p* < 0.05.

Why do networks with longer inhibitory connections have lower inhibitory weights? To answer this, we turned to an analysis of individual synaptic weights within networks, correlating inhibitory synaptic weight against the distance between pre- and postsynaptic neurons. As shown in the example in Figure 7E, longer-range inhibitory synapses tend to be weaker than short-range synapses. This holds true across networks in most of the parameter space of *r*_*inh*_ and *p*_*inh*_ (Figure 7F), with an average correlation coefficient of *r* = −0.32 ± 0.18. The effect is naturally weaker where *r*_*inh*_ is so small that there are few inhibitory connections and little variation in their distance to begin with, but very consistent in networks with *r*_*inh*_ ≥ 2 mm, where the correlation is very strong on average (*r* = −0.43 ± 0.18).

A small caveat is warranted here. Due to our choice of a hard-bounds STDP rule, weights developed into a bimodal distribution with values of either nearly 0 or nearly 1 as noted above and shown in Supplementary Figure 2. Consequently, the observed lower inhibitory weight average for networks with more long-range connections (Figure 7C) and the negative correlations between length and weight within networks (Figure 7F) should be interpreted in the sense that longer-range connections are less likely to be potentiated into an active state than short-range ones. Put differently, networks with longer average connection lengths have a smaller pool of functional (short-range) inhibitory synapses, and thus a lower effective inhibitory degree. Together with the finding of relatively tight E/I balance constraints on criticality (Figure 5), this neatly explains the higher values of Δ*p* in longer-range networks (Figure 7A).

### Long-range synapses experience delayed spike synchrony

To understand why long-range connections fail to potentiate over development, we then considered the underlying learning rule, the Mexican hat shaped inhibitory STDP rule, and its driving factor, the coincidence of pre- and postsynaptic spikes. If long-range connections are not being strengthened, it must be the case that there are few coincident spike pairs relative to non-coincident spike pairs. To confirm this and understand how distance affects spike coincidence, we turned to a sample network that reaches criticality (*r*_*inh*_ = 4 mm, *p*_*inh*_ = 0.7, shown also in Figure 3 and Figure 7). We separated synapses into groups by their distance, then considered all spike pairs with a temporal separation at the synapse of less than 50 ms, plotting their occurrence over developmental time as a heatmap (Figure 8A).

**Figure 8.**
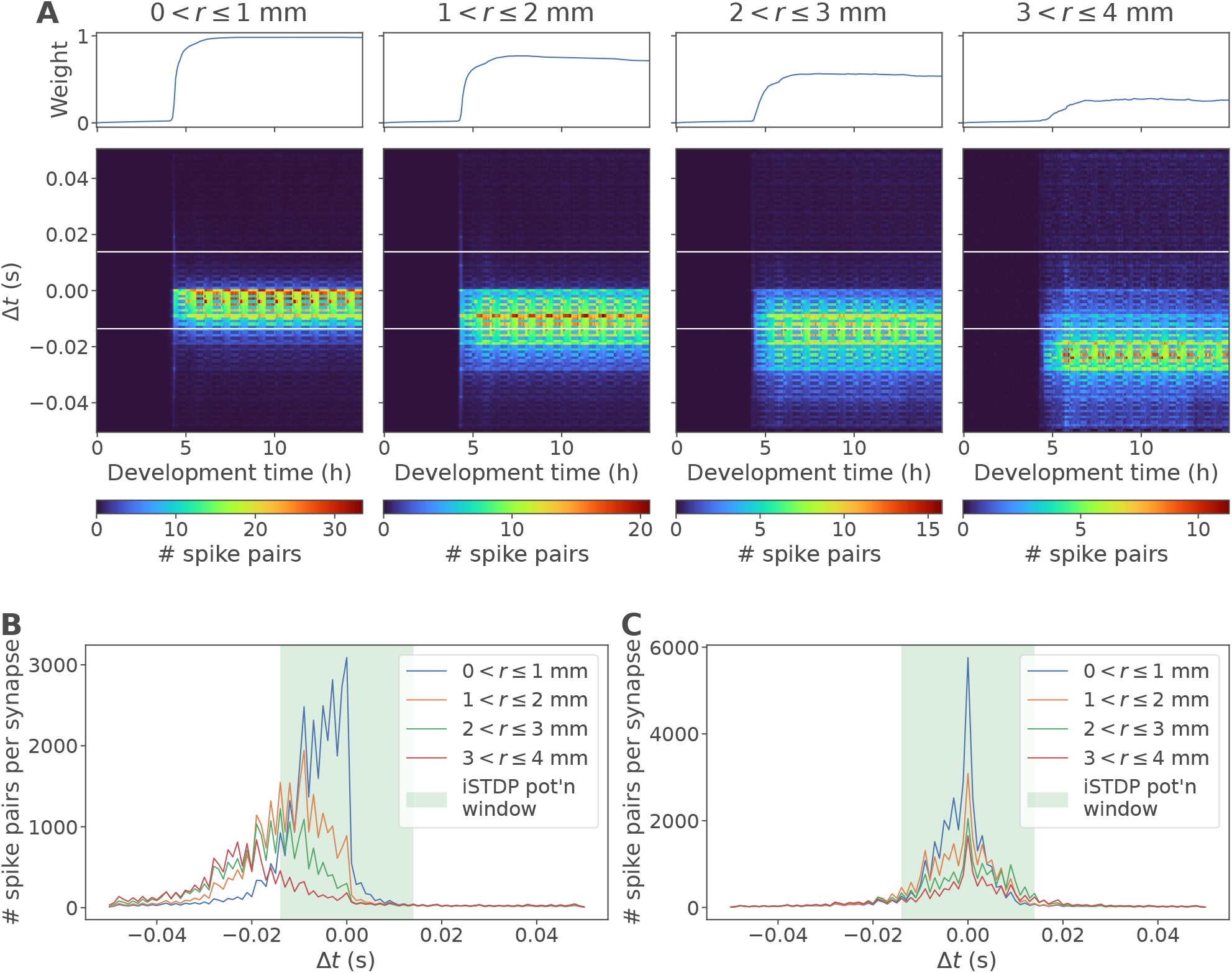
Long-range synapses experience delayed spike synchrony. **(A)** Spike coincidence plots for inhibitory synapses in a sample network (same as Figure 3C and Figure 7E-F), sorted by connection length from the shortest connections (left) to the longest connections (right). The top panel shows the average weight of the connections over development, while the bottom panel shows the density of spikes paired within ±50 ms at their respective *Δt* = *t*_*post*_ − *t*_*pre*_. The white horizontal lines indicate the boundaries of the potentiation window of the inhibitory STDP rule. Spike timing is measured locally at the synapse, mirroring the STDP weight update procedure. **(B)** The same data summed over development time and plotted as lines. The shaded area indicates the potentiation window of the inhibitory STDP rule. **(C)** The same data but using the global timing of spikes instead of the timing local to each synapse.

In all synapse groups, the early developmental period was marked by very low levels of coincident activity, consistent with low overall activity and a lack of growth in inhibitory weights (cf. Figure 3). After the transition to criticality at ∼4.5 hours, all groups showed greatly increased coincident spiking, though with a clear trend towards more frequent coincident spikes in shorter-range synapses (note the different color scales in Figure 8A). Crucially, while the timing of pre- and postsynaptic spikes was approximately synchronous in short-range synapses, long-range synapses showed strong asynchrony, with most spike pairings following a post-before-pre pattern (Δ*t* < 0). This is shown more clearly still in the histogram over all time in Figure 8B.

To understand this asynchrony, consider that the timing of spike pairs in the above was measured at the synapse, which was modelled at the location of the postsynaptic neuron. Therefore, the presynaptic spike timing included a distance-dependent conduction delay, while the postsynaptic spike was recorded at the time of its emission without delay. Correcting for this delay, i.e., recording as the Δ*t* of a spike pair the time difference of the emission of spikes at the respective neuron rather than that of their arrival at the synapse, we see that all spike pairs, from the shortest-range to the longest-range synapses, strongly tend towards synchrony on average (Figure 8C and Supplementary Figure 9). Together with the axonal (i.e., presynaptic) delay and the shape of the inhibitory learning rule, this synchrony in spiking therefore leads to potentiation in short-range synapses, but depression in long-range synapses.

Consequently, many long-range inhibitory synapses are ineffective with weights close to 0, and therefore, networks with a substantial fraction of long-range inhibitory synapses experience both less inhibition and thus a greater tendency towards supercritical dynamics than networks where those same synapses are concentrated in a smaller radius. Tying this back to our connectivity constraints, we should therefore expect networks with small radius *r*_*inh*_ and large connection probability *p*_*inh*_ to show fewer large avalanches than corresponding networks with larger *r*_*inh*_ and smaller *p*_*inh*_. Looking back at Figure 4C, we see exactly this: Networks with *r*_*inh*_ = 2 mm and *p*_*inh*_ = 0.7 (center column, second from the top; mean *k*_*inh*_ = 41, mean Δ*p* = 0.11) are much more likely to be subcritical than networks with *r*_*inh*_ = 3 mm and *p*_*inh*_ = 0.5 (middle row, second from the right; mean *k*_*inh*_ = 45, mean Δ*p* = 0.3), even though the latter have more inhibitory connections.

## Discussion

In this work, we demonstrated that criticality is affected by the distance between inhibitory neurons and their postsynaptic targets, in that long-range inhibitory connections tend to be weaker than short-range connections. Networks with preferentially short-range inhibitory connections are therefore more strongly inhibited, biasing the dynamics towards the subcritical phase. We further showed that synaptic delay is the main cause for the lower weights in long-range synapses, highlighting the role of conduction delays in establishing and maintaining criticality.

While we have explained our findings in terms of effective degree as a result of the strongly bimodal weight distributions observed, related work has employed both hard-bound STDP rules leading to bimodal weight distributions and soft-bound STDP rules leading to more realistic unimodal weight distributions but found no substantial difference in the resulting patterns of activity and criticality (Rubinov et al. 2011). This allows us to speculate that in our case, too, a soft-bound rule would lead to similar outcomes in terms of activity while leading to explanations in terms of graded variations of synaptic weight.

Strictly speaking, our model does not demonstrate self-organized criticality, as shown by the narrow band of parameter settings that lead to criticality, and consequently the large number of networks with persistently sub- or supercritical dynamics. This stands in contrast to related work, where short-term plasticity (Zeraati, Priesemann, and Levina 2021) or long-term plasticity with high learning rates (Rubinov et al. 2011) directly contribute to the establishment and maintenance of criticality in a comparatively broad region of parameter space. With our approach to plasticity, with low learning rates enforcing a scale separation between development and dynamics, we chose to highlight the structural, rather than dynamic, contributions to criticality.

There is a clear connection between our results and cortical connectivity patterns, which are widely known to strongly favor short-range local connections for inhibitory synapses (Fino, Packer, and Yuste 2013). While this has been attributed in part to a reduction of wiring cost (Samu, Seth, and Nowotny 2014; Buzsáki et al. 2004), our work suggests that short-range connectivity also emerges as a consequence of an interplay between the homeostatic quasi-Hebbian learning rule thought to dominate inhibitory long-term plasticity (Vogels et al. 2011) and the frequent occurrence of neuronal avalanches in near-critical networks. In our model, long-range synapses are not weakened because distant neurons are not acting in synchrony – indeed, on average, they are – but rather because transmission delays hinder the consistent temporal coordination of pre- and postsynaptic events. However, unlike our model, cortex possesses higher-order structure, which may bias activity propagation in ways that allow coordinated firing, and therefore strong inhibitory connectivity, over longer distances along frequently used signaling pathways.

Related work has shown that higher-order structure in the form of hierarchical modularity is conducive to critical dynamics (Wang and Zhou 2012; Okujeni and Egert 2023). In some cases, even modularity in only the inhibitory connections is sufficient to promote criticality, while excitatory modularity only plays a secondary role (Rubinov et al. 2011). This aligns well with our results, since neural modules or clusters are likely to fire in a coordinated manner, supporting the formation of strong inhibitory synapses. How such modules form in the first place, and what defines their boundaries, however, is outside the scope of our exploration.

A key limitation in our model is its small size, corresponding to just a single module in the sense of the larger, hierarchically structured Rubinov et al. (2011) model. It is arguably difficult to draw conclusions about criticality in a system of only 100 neurons. However, as demonstrated, it is clearly possible to smoothly interpolate our model from sub-to supercritical states with minor parameter changes, indicating that the power law scaling observed in some networks does in fact derive from criticality, not from some other process (Beggs and Timme 2012).

Whilst we showed that transmission delays have a notable impact on the (effective) structure of a spatially defined network, we have not investigated their dynamical implications. Theoretical considerations have shown that transmission delays affect the time scale of relaxation towards steady state activity when an external input is applied but not the steady state activity itself (Larremore et al. 2011). In networks of finite size, we suspect that the dynamics may well depend on transmission delays in nontrivial ways and would encourage further research along these lines.

## Author Contributions

FBK: Investigation, Methodology, Software, Visualization, Writing – original draft, Writing – review and editing; TD: Formal analysis, Investigation, Methodology, Writing – review and editing; ZCC: Conceptualization, Funding acquisition, Supervision, Writing – review and editing.

## Acknowledgments

We thank Drs Hirokazu Takahashi and Narumitsu Ikeda for their insightful discussion and inspiration that shaped the model design.

## Data Availability Statement

The code used to generate all data analyzed in this study and its figures, as well as the partially processed data underlying the figures and quantitative information presented herein, can be found at https://github.com/kernfel/inhibitory-constraints-criticality.

## References

Bak, Per, Chao Tang, and Kurt Wiesenfeld. 1987. “Self-Organized Criticality: An Explanation of the 1/f Noise.” Physical Review Letters 59 (4): 381–84. 10.1103/PhysRevLett.59.381.

Beggs, John M. 2007. “The Criticality Hypothesis: How Local Cortical Networks Might Optimize Information Processing.” Philosophical Transactions of the Royal Society A: Mathematical, Physical and Engineering Sciences 366 (1864): 329–43. 10.1098/rsta.2007.2092.

Beggs, John M., and Dietmar Plenz. 2003. “Neuronal Avalanches in Neocortical Circuits.” Journal of Neuroscience 23 (35): 11167–77. 10.1523/JNEUROSCI.23-35-11167.2003.

Beggs, John M., and Dietmar Plenz. 2004. “Neuronal Avalanches Are Diverse and Precise Activity Patterns That Are Stable for Many Hours in Cortical Slice Cultures.” Journal of Neuroscience 24 (22): 5216–29. 10.1523/JNEUROSCI.0540-04.2004.

Beggs, John M., and Nicholas Timme. 2012. “Being Critical of Criticality in the Brain.” Frontiers in Physiology 3 (June):163. 10.3389/fphys.2012.00163.

Benayoun, Marc, Jack D. Cowan, Wim van Drongelen, and Edward Wallace. 2010. “Avalanches in a Stochastic Model of Spiking Neurons.” PLOS Computational Biology 6 (7): e1000846. 10.1371/journal.pcbi.1000846.

Bertschinger, Nils, and Thomas Natschläger. 2004. “Real-Time Computation at the Edge of Chaos in Recurrent Neural Networks.” Neural Computation 16 (7): 1413–36. 10.1162/089976604323057443.

Buzsáki, György, Caroline Geisler, Darrell A. Henze, and Xiao-Jing Wang. 2004. “Interneuron Diversity Series: Circuit Complexity and Axon Wiring Economy of Cortical Interneurons.” Trends in Neurosciences 27 (4): 186–93. 10.1016/j.tins.2004.02.007.

Del Papa, Bruno, Viola Priesemann, and Jochen Triesch. 2017. “Criticality Meets Learning: Criticality Signatures in a Self-Organizing Recurrent Neural Network.” PLOS ONE 12 (5): e0178683. 10.1371/journal.pone.0178683.

Fino, Elodie, Adam M. Packer, and Rafael Yuste. 2013. “The Logic of Inhibitory Connectivity in the Neocortex.” The Neuroscientist 19 (3): 228–37. 10.1177/1073858412456743.

Haldeman, Clayton, and John M. Beggs. 2005. “Critical Branching Captures Activity in Living Neural Networks and Maximizes the Number of Metastable States.” Physical Review Letters 94 (5): 058101. 10.1103/PhysRevLett.94.058101.

Hesse, Janina, and Thilo Gross. 2014. “Self-Organized Criticality as a Fundamental Property of Neural Systems.” Frontiers in Systems Neuroscience 8 (September). 10.3389/fnsys.2014.00166.

Ikeda, Narumitsu, Dai Akita, and Hirokazu Takahashi. 2023. “Noise and Spike-Time-Dependent Plasticity Drive Self-Organized Criticality in Spiking Neural Network: Toward Neuromorphic Computing.” Applied Physics Letters 123 (2): 023701. 10.1063/5.0152633.

Kinouchi, Osame, and Mauro Copelli. 2006. “Optimal Dynamical Range of Excitable Networks at Criticality.” Nature Physics 2 (5): 348–51. 10.1038/nphys289.

Larremore, Daniel B., Woodrow L. Shew, Edward Ott, and Juan G. Restrepo. 2011. “Effects of Network Topology, Transmission Delays, and Refractoriness on the Response of Coupled Excitable Systems to a Stochastic Stimulus.” Chaos: An Interdisciplinary Journal of Nonlinear Science 21 (2): 025117. 10.1063/1.3600760.

Legenstein, Robert, and Wolfgang Maass. 2007. “Edge of Chaos and Prediction of Computational Performance for Neural Circuit Models.” Neural Networks, Echo State Networks and Liquid State Machines, 20 (3): 323–34. 10.1016/j.neunet.2007.04.017.

Mazzoni, Alberto, Frédéric D. Broccard, Elizabeth Garcia-Perez, Paolo Bonifazi, Maria Elisabetta Ruaro, and Vincent Torre. 2007. “On the Dynamics of the Spontaneous Activity in Neuronal Networks.” PLOS ONE 2 (5): e439. 10.1371/journal.pone.0000439.

Okujeni, Samora, and Ulrich Egert. 2023. “Structural Modularity Tunes Mesoscale Criticality in Biological Neuronal Networks.” Journal of Neuroscience 43 (14): 2515–26. 10.1523/JNEUROSCI.1420-22.2023.

Plenz, Dietmar, Tiago L. Ribeiro, Stephanie R. Miller, Patrick A. Kells, Ali Vakili, and Elliott L. Capek. 2021. “Self-Organized Criticality in the Brain.” Frontiers in Physics 9 (July). 10.3389/fphy.2021.639389.

Poil, Simon-Shlomo, Richard Hardstone, Huibert D. Mansvelder, and Klaus Linkenkaer-Hansen. 2012. “Critical-State Dynamics of Avalanches and Oscillations Jointly Emerge from Balanced Excitation/Inhibition in Neuronal Networks.” Journal of Neuroscience 32 (29): 9817–23. 10.1523/JNEUROSCI.5990-11.2012.

Ponce-Alvarez, Adrián, Adrien Jouary, Martin Privat, Gustavo Deco, and Germán Sumbre. 2018. “Whole-Brain Neuronal Activity Displays Crackling Noise Dynamics.” Neuron 100 (6): 1446–1459.e6. 10.1016/j.neuron.2018.10.045.

Rubinov, Mikail, Olaf Sporns, Jean-Philippe Thivierge, and Michael Breakspear. 2011. “Neurobiologically Realistic Determinants of Self-Organized Criticality in Networks of Spiking Neurons.” PLOS Computational Biology 7 (6): e1002038. 10.1371/journal.pcbi.1002038.

Samu, David, Anil K. Seth, and Thomas Nowotny. 2014. “Influence of Wiring Cost on the Large-Scale Architecture of Human Cortical Connectivity.” PLOS Computational Biology 10 (4): e1003557. 10.1371/journal.pcbi.1003557.

Shew, Woodrow L., Hongdian Yang, Thomas Petermann, Rajarshi Roy, and Dietmar Plenz. 2009. “Neuronal Avalanches Imply Maximum Dynamic Range in Cortical Networks at Criticality.” Journal of Neuroscience 29 (49): 15595–600. 10.1523/JNEUROSCI.3864-09.2009.

Shew, Woodrow L., Hongdian Yang, Shan Yu, Rajarshi Roy, and Dietmar Plenz. 2011. “Information Capacity and Transmission Are Maximized in Balanced Cortical Networks with Neuronal Avalanches.” Journal of Neuroscience 31 (1): 55–63. 10.1523/JNEUROSCI.4637-10.2011.

Song, Sen, Kenneth D. Miller, and L. F. Abbott. 2000. “Competitive Hebbian Learning through Spike-Timing-Dependent Synaptic Plasticity.” Nature Neuroscience 3 (9): 919–26. 10.1038/78829.

Stepp, Nigel, Dietmar Plenz, and Narayan Srinivasa. 2015. “Synaptic Plasticity Enables Adaptive Self-Tuning Critical Networks.” PLOS Computational Biology 11 (1): e1004043. 10.1371/journal.pcbi.1004043.

Tetzlaff, Christian, Samora Okujeni, Ulrich Egert, Florentin Wörgötter, and Markus Butz. 2010. “Self-Organized Criticality in Developing Neuronal Networks.” PLOS Computational Biology 6 (12): e1001013. 10.1371/journal.pcbi.1001013.

Tsodyks, Misha V., and Henry Markram. 1997. “The Neural Code between Neocortical Pyramidal Neurons Depends on Neurotransmitter Release Probability.” Proceedings of the National Academy of Sciences 94 (2): 719–23. 10.1073/pnas.94.2.719.

Vogels, T. P., H. Sprekeler, F. Zenke, C. Clopath, and W. Gerstner. 2011. “Inhibitory Plasticity Balances Excitation and Inhibition in Sensory Pathways and Memory Networks.” Science 334 (6062): 1569–73. 10.1126/science.1211095.

Wang, Sheng-Jun, and Changsong Zhou. 2012. “Hierarchical Modular Structure Enhances the Robustness of Self-Organized Criticality in Neural Networks.” New Journal of Physics 14 (2): 023005. 10.1088/1367-2630/14/2/023005.

Wilkerson, Galen J., and Sotiris Moschoyiannis. 2021. “Logic and Learning in Network Cascades.” Network Science 9 (S1): S157–74. 10.1017/nws.2021.3.

Zeraati, Roxana, Viola Priesemann, and Anna Levina. 2021. “Self-Organization Toward Criticality by Synaptic Plasticity.” Frontiers in Physics 9 (April). 10.3389/fphy.2021.619661.

