## Supplementary Material for "Effects of Spatial Constraints of Inhibitory Connectivity on the Dynamical Development of Criticality in Spiking Networks"

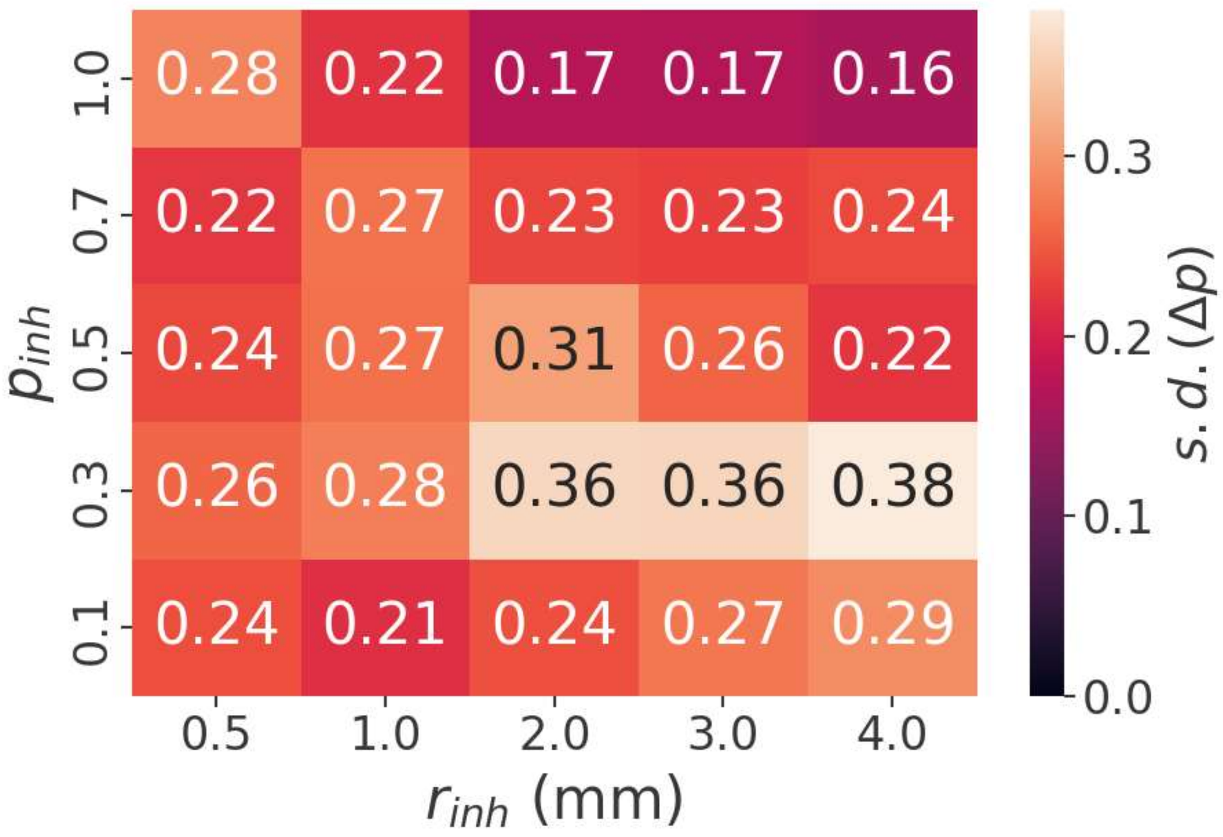


**Supplementary Figure 1.** Standard deviation of $\Delta p$ across the grid of $r_{inh}$ and $p_{inh}$. $\Delta p$ was calculated in each run of each network from activity between 13 and 15 hours. The standard deviation was then calculated across all runs and networks (n=50) in each grid cell.

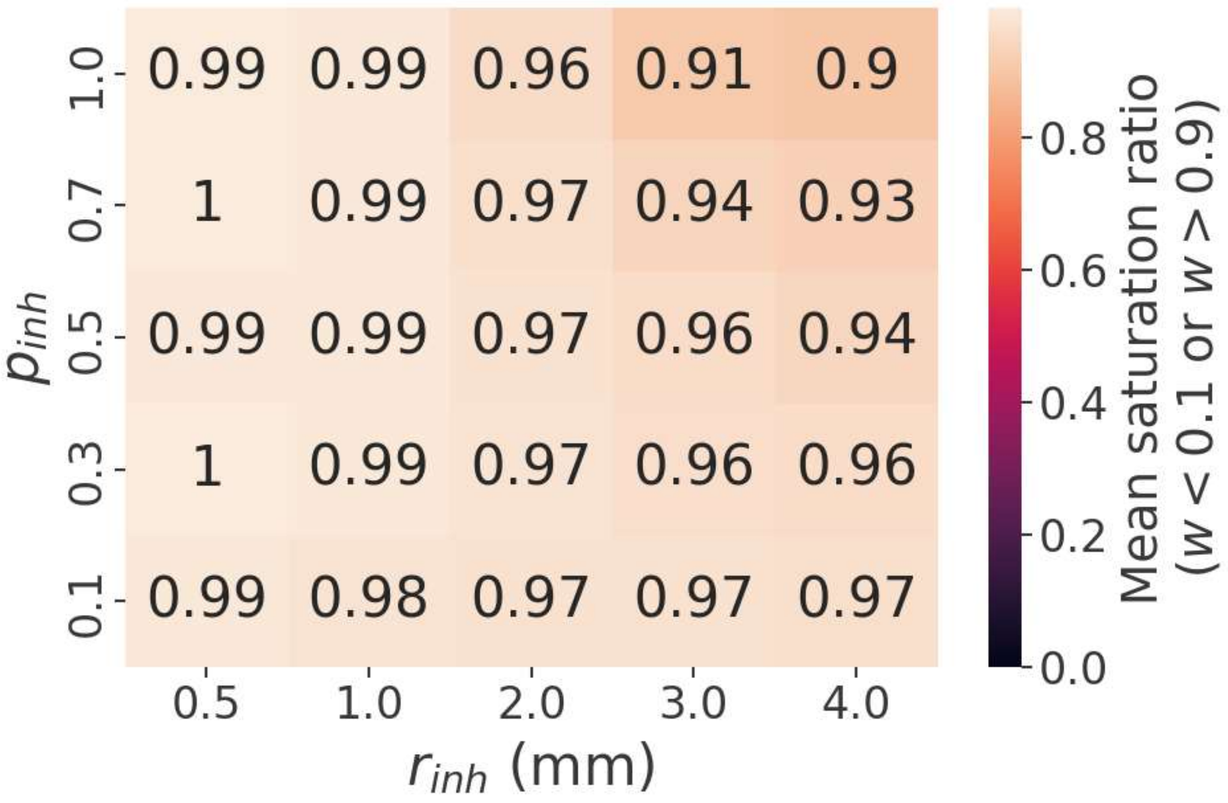


**Supplementary Figure 2.** Saturation ratio of networks, i.e., the fraction of weights within 0.1 of the lower and upper bounds, across the grid of $r_{inh}$ and $p_{inh}$, after development.

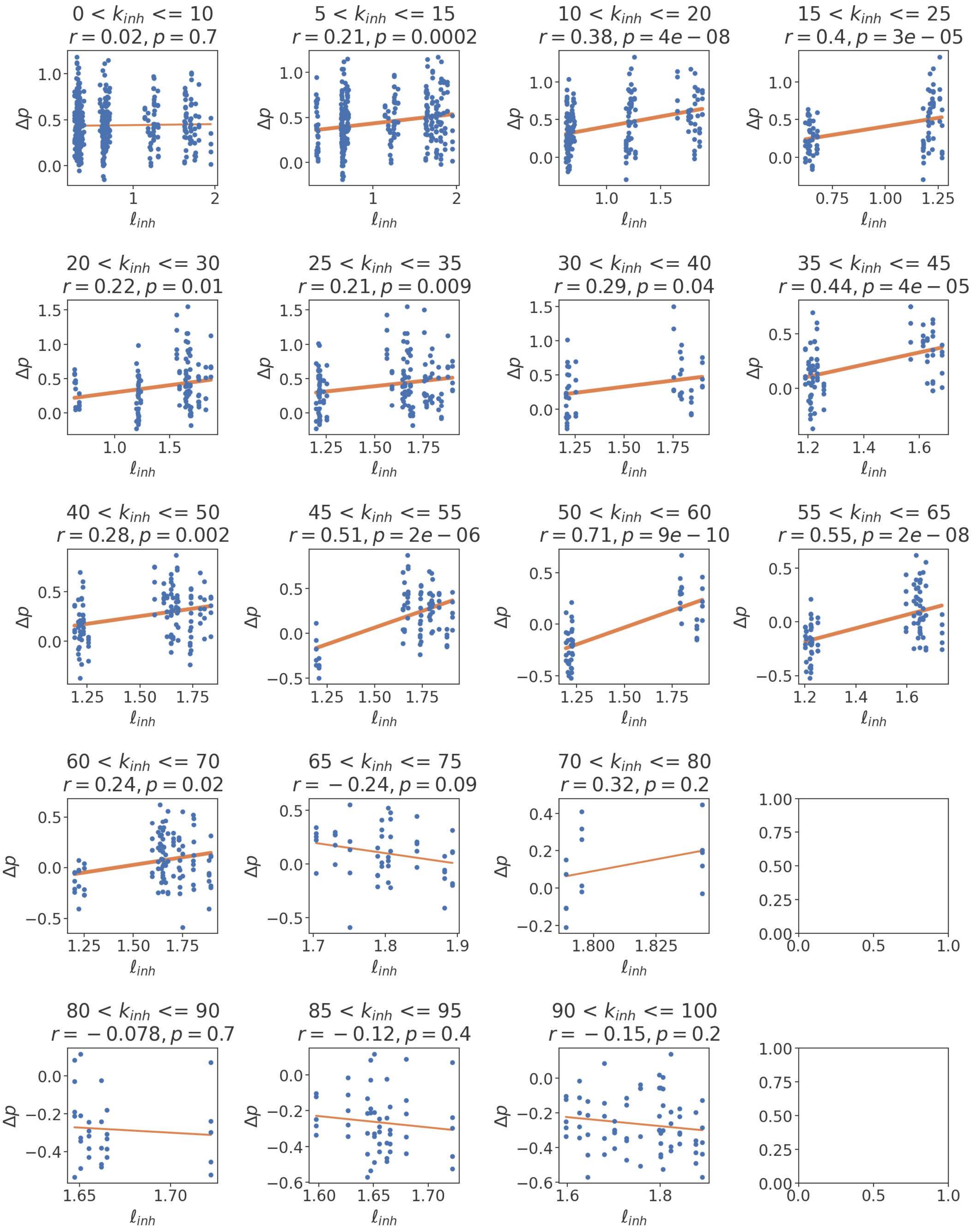


**Supplementary Figure 3.** Scatter plots and associated linear fits relating $l_{inh}$ to $\Delta p$ in each slice of $k_{inh}$, see Figure 6E.

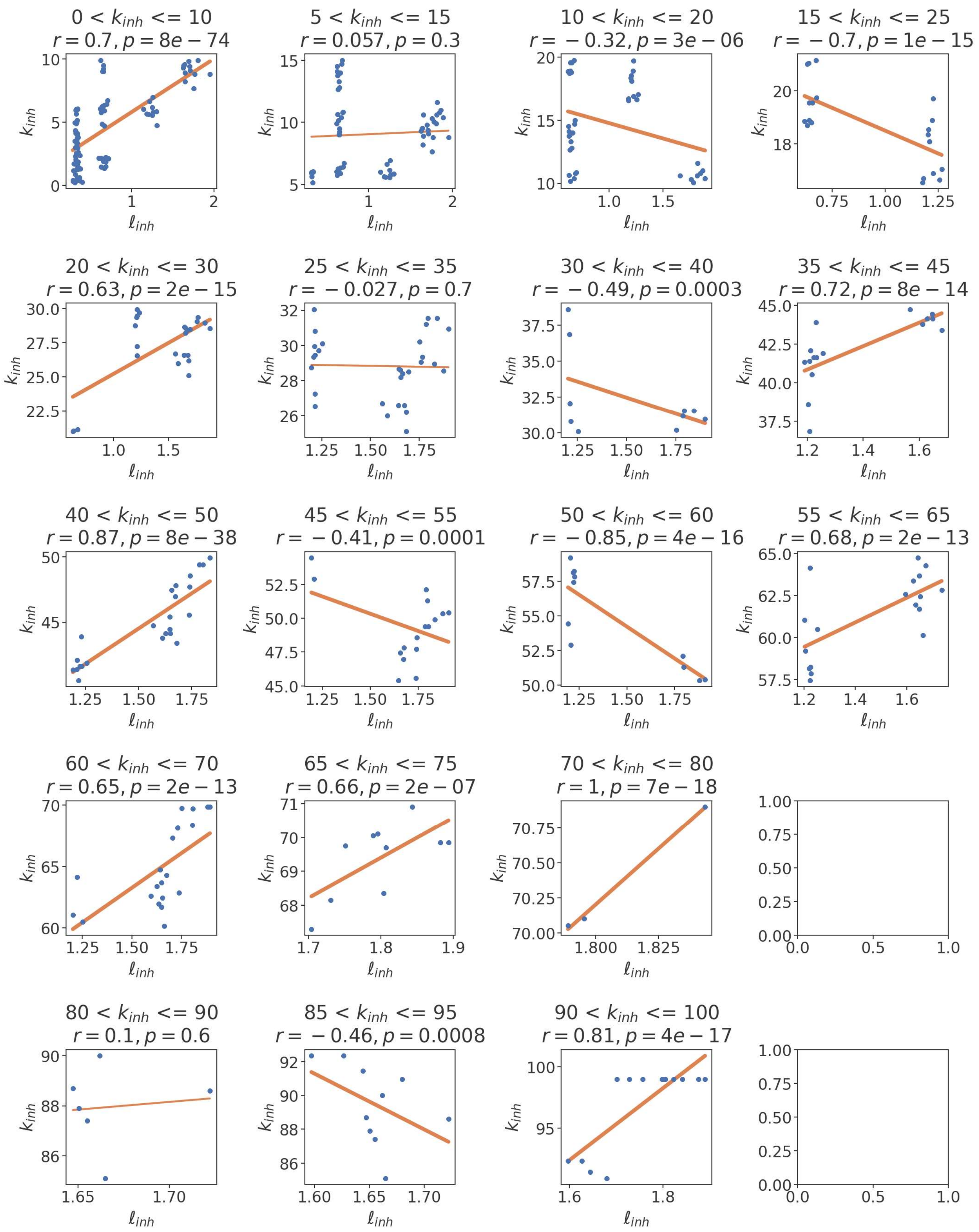


**Supplementary Figure 4.** Scatter plots and associated linear fits relating $l_{inh}$ to $k_{inh}$ in each slice of $k_{inh}$, see Figure 6F.

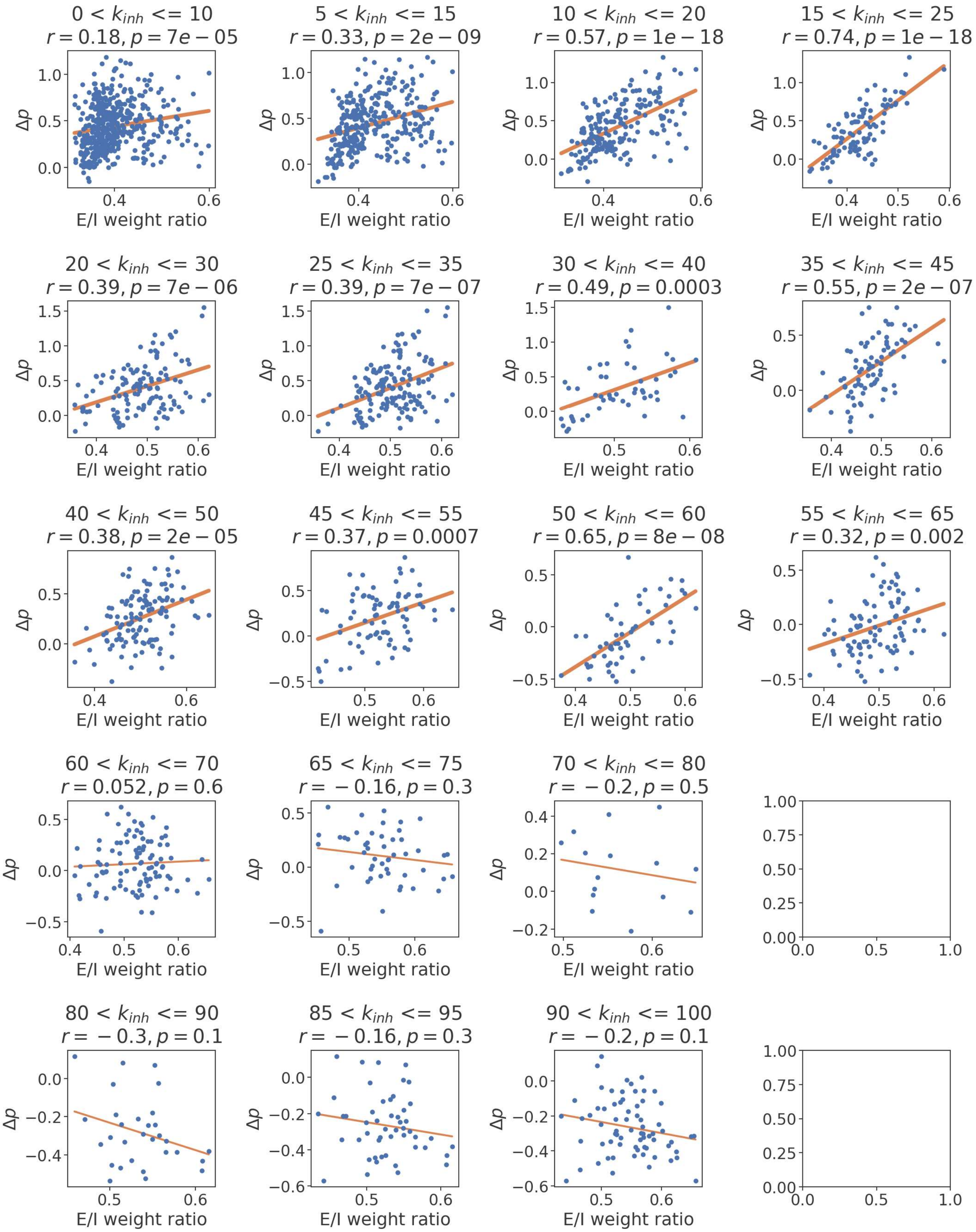


**Supplementary Figure 5.** Scatter plots and associated linear fits relating the E/I ratio to $\Delta p$ in each slice of $k_{inh}$, see Figure 7A.

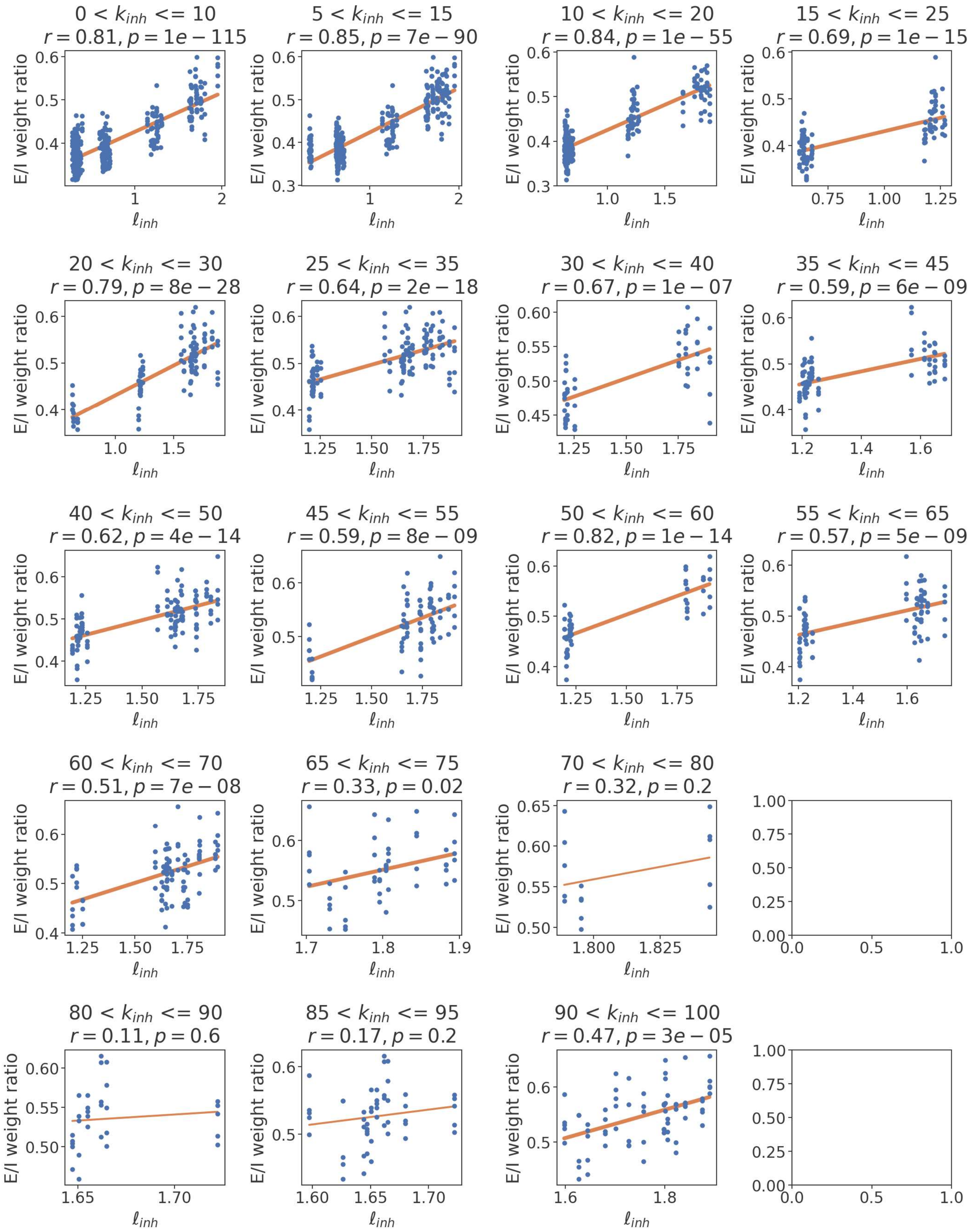


**Supplementary Figure 6.** Scatter plots and associated linear fits relating $l_{inh}$ to the E/I ratio in each slice of $k_{inh}$, see Figure 7B.

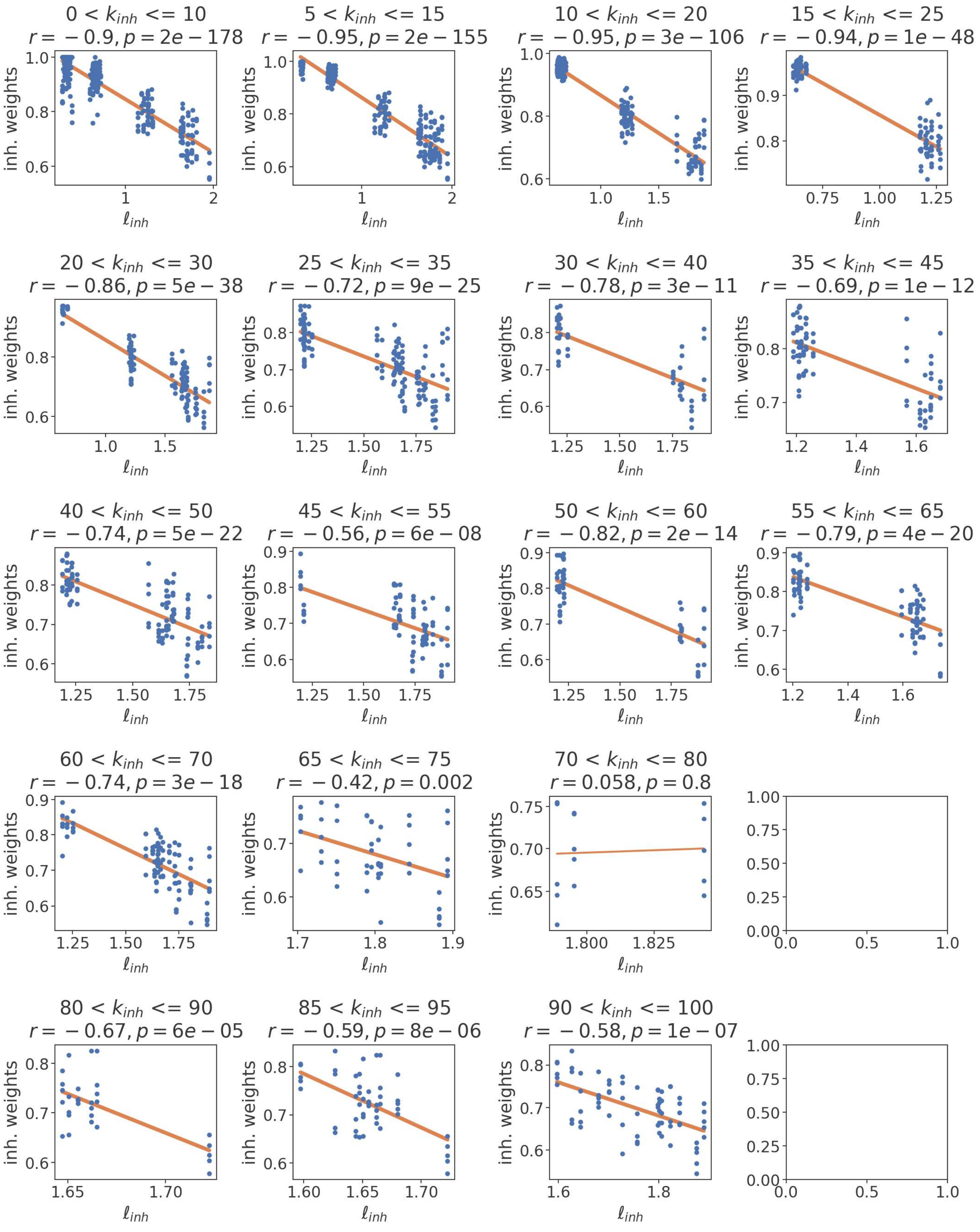


**Supplementary Figure 7.** Scatter plots and associated linear fits relating $l_{inh}$ to the inhibitory weight average in each slice of $k_{inh}$, see Figure 7C.

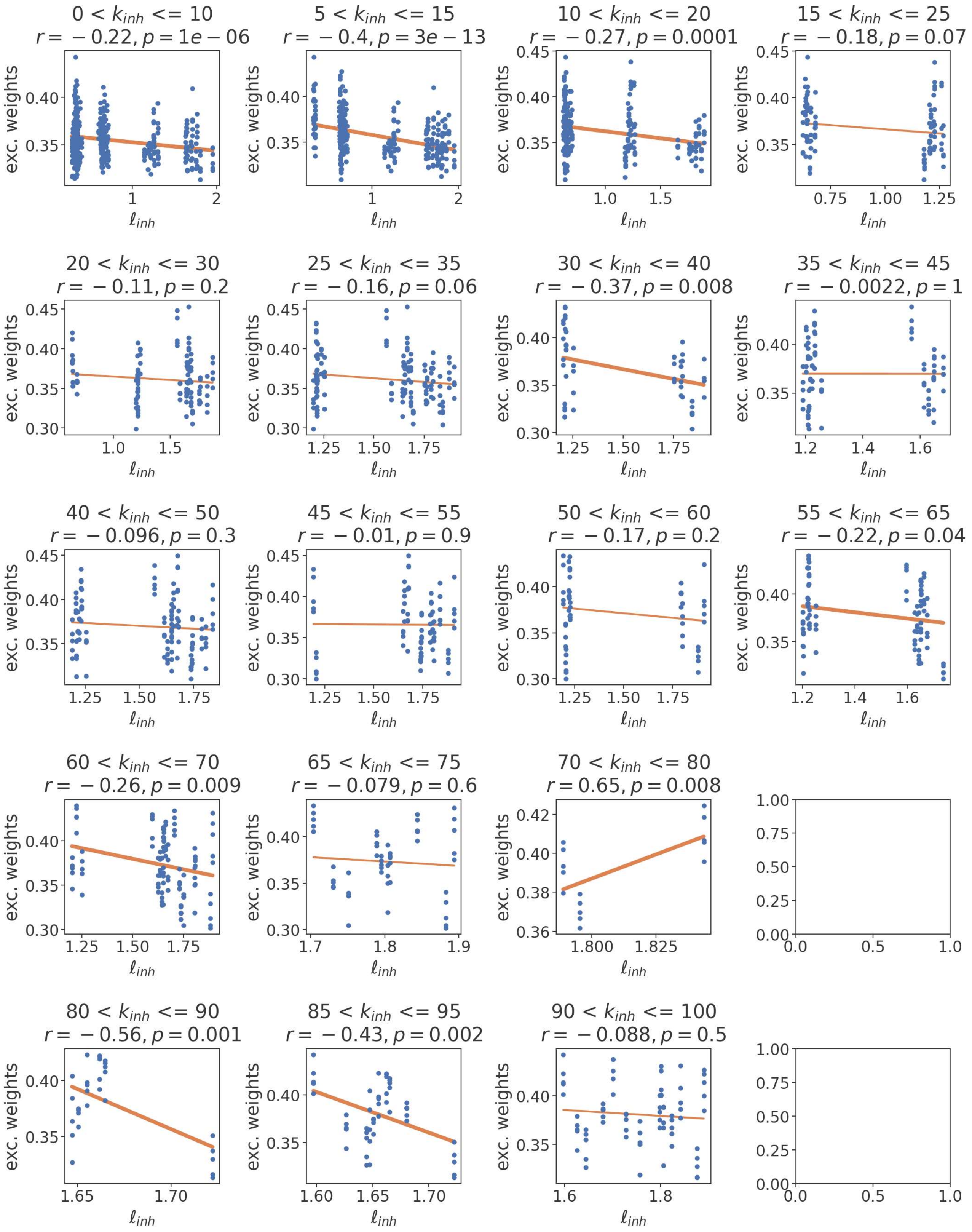


**Supplementary Figure 8.** Scatter plots and associated linear fits relating $l_{inh}$ to the excitatory weight average in each slice of $k_{inh}$, see Figure 7D.

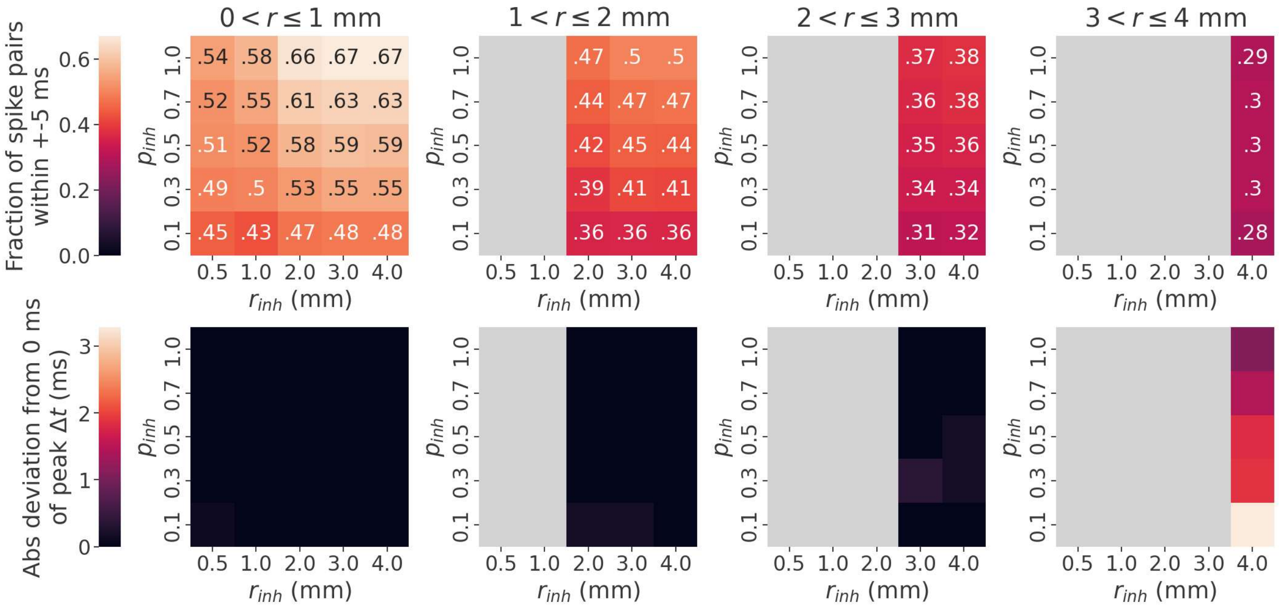


**Supplementary Figure 9.** Synchronization indicators in inhibitory synapses of various lengths across all networks. Top, fraction of spike pairs occurring within +-5 ms; bottom, deviation of the most frequent pre/post spike time difference from 0 ms. Indicators were calculated within each network separately, then averaged across all 50 networks in each grid cell. Spike timings were recorded from all spike pairs occurring within +-50 ms and calculated without including conduction delays, as in Figure 8C. Grey areas indicate no data (i.e., there were no long-range synapses in short-range parameter settings).
